# Taxonomic and functional concordance between full-length ONT 16S and ONT shotgun metagenomics in the canine gut microbiome

**DOI:** 10.64898/2026.09.26.754683

**Authors:** Natheer Jameel Yaseen, Balázs Kakuk, Gábor Gulyás, Md Asaduzzaman, Zsolt Boldogkői, Dóra Tombácz

## Abstract

**Background:** Full-length Oxford Nanopore Technologies (ONT) 16S rRNA sequencing provides a scalable view of microbial community composition and can support phylogeny-based functional prediction, but it is not equivalent to shotgun metagenomics. We asked which biological conclusions are preserved when the same canine fecal specimens are profiled by full-length ONT 16S and ONT whole-genome shotgun (WGS) sequencing, and how their agreement depends on analytical scale, reference representation and classifier.

**Methods:** Ninety-seven fecal specimens from 51 dogs were profiled with both assays from the same DNA extract. Functional profiles predicted from NanoASV/NanoPredict with PICRUSt2 were compared with WGS-supported KEGG Ortholog (KO) profiles generated by Kadath. Taxonomy was benchmarked in a source-genome-matched RefSeq universe and in a host-specific DogMAG universe using minitax and Kraken2. Agreement was evaluated at whole-profile, feature-abundance, detection, between-sample structure and biological-inference scales. Age-associated transfer was assessed with dog-aware continuous mixed models, grouped signed-score analyses and paired/dog-blocked PERMANOVA.

**Results:** Functional whole-profile concordance was high: median within-sample CLR Spearman correlations ranged from 0.781 to 0.860 across developmental strata, while between-sample functional structure remained significant by Mantel (rho=0.543) and Procrustes (r=0.693; both p=0.001). Feature-wise transfer was substantially weaker (median KO-wise CLR Spearman=0.318). Continuous age-associated KO slopes showed substantial cross-assay concordance (Spearman=0.727; signed-score Spearman=0.753; direction agreement=77.9%), although 1,290/5,258 eligible KOs retained significant assay-by-age interactions. Taxonomically, exact genus/species abundance agreement was much lower than agreement in between-sample ecological structure. Host-specific DogMAG improved species-level median Spearman from 0.261 to 0.656 for minitax SpeciesEstimate and from 0.181 to 0.512 for Kraken2. The classifier effect was independent of reference choice: under both RefSeq and DogMAG, minitax yielded stronger 16S–WGS concordance than Kraken2, with all eight prespecified RefSeq paired genus/species endpoints and all 10 DogMAG primary paired endpoints significant after BH correction. The same ordering extended to developmental inference, with DogMAG genus/species age-slope concordance of 0.795/0.799 for SpeciesEstimate versus 0.693/0.702 for Kraken2. Taxonomic Aitchison PERMANOVA detected age-associated structure in every assay/reference/classifier/rank combination, whereas age-by-assay interactions were consistently significant but small (R2 approximately 1.1–2.2%). Stricter NanoASV identity thresholds removed substantial 16S abundance without improving species-level agreement.

**Conclusions:** The extent of cross-assay agreement depends on the level of analysis. Full-length ONT 16S preserves broad functional organization, ecological structure and much of the direction of age-associated change, but exact fine-rank composition, individual-feature abundance and effect magnitude remain assay dependent. Host-specific reference representation substantially narrows the taxonomic gap, and classifier choice exerts an additional independent effect: within the same matched reference set, minitax consistently yields stronger 16S–WGS concordance than Kraken2 across abundance, detection, ecological-distance and developmental-inference endpoints. Full-length ONT 16S is therefore well suited to broad ecological screening and hypothesis generation, whereas WGS remains preferable when conclusions depend on quantitative fine-rank composition, directly supported gene content or precise feature-level effect estimates.

## Introduction

Marker-gene and shotgun metagenomic sequencing describe overlapping but non-equivalent aspects of microbial communities. Full-length 16S rRNA sequencing can be applied economically to large longitudinal cohorts and provides substantially more phylogenetic information than short hypervariable-region amplicons, but it still observes a single multicopy marker rather than complete genomes. Shotgun metagenomics samples community genomic content directly and can support gene-level annotation and finer taxonomic assignment, although its output remains dependent on sequencing depth, assembly, reference coverage and classifier behavior. A recent critical review of head-to-head benchmarking emphasized that sequencing strategy should be treated as a study-design choice rather than a neutral technical substitution, and also highlighted the limited direct benchmarking evidence connecting current full-length ONT 16S workflows with long-read shotgun metagenomics (Albastaki and Smith 2026). Paired comparisons can establish which biological conclusions are stable across assays and how analytical choices affect that stability.

ONT full-length 16S sequencing has improved taxonomic resolution in several settings (Cuscó et al. 2018; Curry et al. 2022; Ni et al. 2023), and longitudinal canine ONT 16S data recover structured variation associated with host development and husbandry (Asaduzzaman et al. 2026). Paired Illumina short-read shotgun metagenomic and full-length ONT 16S profiling of Pumi puppies further showed that early-life gut maturation follows a shared age–diet trajectory within persistent host-specific structure (Járay et al. 2026). Nevertheless, fine-rank abundance estimates remain sensitive to amplification, rRNA operon copy number, within-lineage 16S similarity, sequencing error and reference composition. A recent canine comparison of 16S workflows similarly showed that broad community patterns can be shared while taxon-level abundance and age-associated results remain workflow dependent (Polacchini et al. 2026). Our multi-platform canine dataset likewise showed that DNA extraction, primer configuration and sequencing platform each shape canine fecal microbiome profiles (Kakuk et al. 2026a), and a cross-comparison of metagenomic profiling strategies introduced minitax as a classifier that gives consistent results across platforms and methods (Gulyás et al. 2024). WGS avoids marker-gene amplification but introduces its own dependencies, including genome representation, read assignment, genome size and the handling of unclassified sequence. For paired-assay benchmarking, WGS is therefore best treated as a more direct operational comparator rather than as biological ground truth.

Reference compatibility and classifier behavior are distinct sources of taxonomic disagreement. A targeted-loci 16S collection and a WGS genome database may use the same taxonomic names while representing different organism sets, loci and resolvable lineages. Harmonizing names cannot recover taxa absent from one reference set or make a conserved 16S sequence uniquely identify a genome. Conversely, even when the reference set is held constant, classifiers can distribute ambiguous reads differently because their search strategies, evidence aggregation and abundance estimators differ. This distinction is especially relevant in canine microbiome research because dog-specific genome catalogs recover substantial host-associated sequence space that is incompletely represented in generic public resources (Castillo-Fernandez et al. 2026). The dog is also an increasingly relevant translational model for microbiome-associated health and aging (Boldogkői and Tombácz 2026), and dog-wise long-read assemblies now provide a host-specific genome resource, DogMAG, for canine gut metagenomics (Kakuk et al. 2026b). Comparing classifiers within a shared reference allows their effects to be assessed separately from those of reference composition.

Functional agreement also depends on the scale of comparison. PICRUSt2 infers expected gene content from phylogenetic placement of marker-gene sequences (Douglas et al. 2020), whereas WGS can support genes directly from sample sequence. Broad functional profiles may agree because many core functions are stable and widespread even when individual gene families, strain-specific functions or host-associated differential signals are poorly reproduced. Previous matched-assay studies have therefore emphasized that high global correlation does not guarantee transfer of differential or metadata-associated inference (Sun, Jones, and Fodor 2020; Matchado et al. 2024). A previous study compared ONT full-length 16S with Ion Torrent shallow shotgun metagenomics in 43 human stool samples and found method-dependent taxon detection and beta-diversity differences, but the two assays were classified against different reference systems and the study did not evaluate long-read WGS, reference harmonization, predicted-versus-supported functional profiles, or transfer of longitudinal metadata-associated effects (Chitcharoen et al. 2025). The same logic applies to taxonomy: preservation of between-sample community structure does not imply equality of species proportions, and similar abundance profiles do not guarantee that the same taxa will carry the same host-associated signal.

Here we compared matched full-length ONT V1–V9 16S and ONT WGS profiles from 97 canine fecal specimens at several levels of analysis. To our knowledge, no previous matched-sample study has combined full-length ONT 16S with long-read ONT shotgun metagenomics while simultaneously controlling the taxonomic reference set, comparing classifier behavior within that reference, benchmarking predicted versus WGS-supported function, and testing repeated-measures metadata-associated inference transfer. Functional analyses separated whole-profile similarity, individual KO trajectories, multivariate community-level inference and feature-level age-associated effects. Taxonomic analyses separated within-sample abundance agreement, detection overlap, between-sample ecological structure, multivariate age structure and taxon-specific age-effect transfer. We first used a source-genome-matched RefSeq benchmark to hold the source genomes constant and then repeated the fine-rank comparison using the host-specific DogMAG reference. Because minitax and Kraken2 were run on the same specimens and references, reference effects could be distinguished from classifier effects. Shared-feature restrictions, abundance-scaling sensitivities and read-count-weighted NanoASV reassignment were used to examine residual disagreement. These analyses assessed which biological conclusions were reproducible across assays and whether analytical adjustments reduced the remaining differences.

## Materials and methods

### Study design, specimen pairing and sequencing

The analysis used a set of 97 canine fecal specimens for which ONT full-length V1–V9 16S and ONT WGS data could be reconciled at the specimen level. The 16S profiles are a matched subset of the longitudinal all-kennel canine dataset reported by Asaduzzaman et al. (Asaduzzaman et al. 2026). The set represents 51 verified dogs from six kennels and was defined by exact dog/date matching, library-level quality review and reconciliation of duplicate or resequenced WGS libraries. Verified dog identity was used as the grouping unit in all repeated-measures analyses.

The WGS data were generated in two acquisition branches. The original WGS collection was sequenced on ONT PromethION, whereas additional libraries were sequenced on ONT MinION. After specimen reconciliation, the 97-pair set comprised 62 specimens represented by PromethION and 35 by MinION; an added library corresponding to already represented biology was not counted as an additional specimen. The extraction, native-barcoding, ligation-library and Dorado super-accurate basecalling framework was otherwise kept consistent across the WGS branches.

Age was recorded for each specimen as biological age in weeks. The primary categorical strata were neonatal/early (<3.5 weeks), weaning (3.5–<8 weeks), post-weaning/juvenile (8–<52 weeks) and adult (>=52 weeks), containing 12, 11, 24 and 50 specimens, respectively. For continuous developmental models, the primary covariate was standardized `log1p(age_weeks)` because the cohort spans neonatal development through late adulthood; standardized raw age in weeks was retained as a sensitivity. Because age, kennel and WGS acquisition branch are not fully balanced, age-associated analyses are interpreted primarily as cross-method transfer tests rather than independent proof of developmental causality.

The analytical design is summarized in Figure 1.

**Figure 1:**
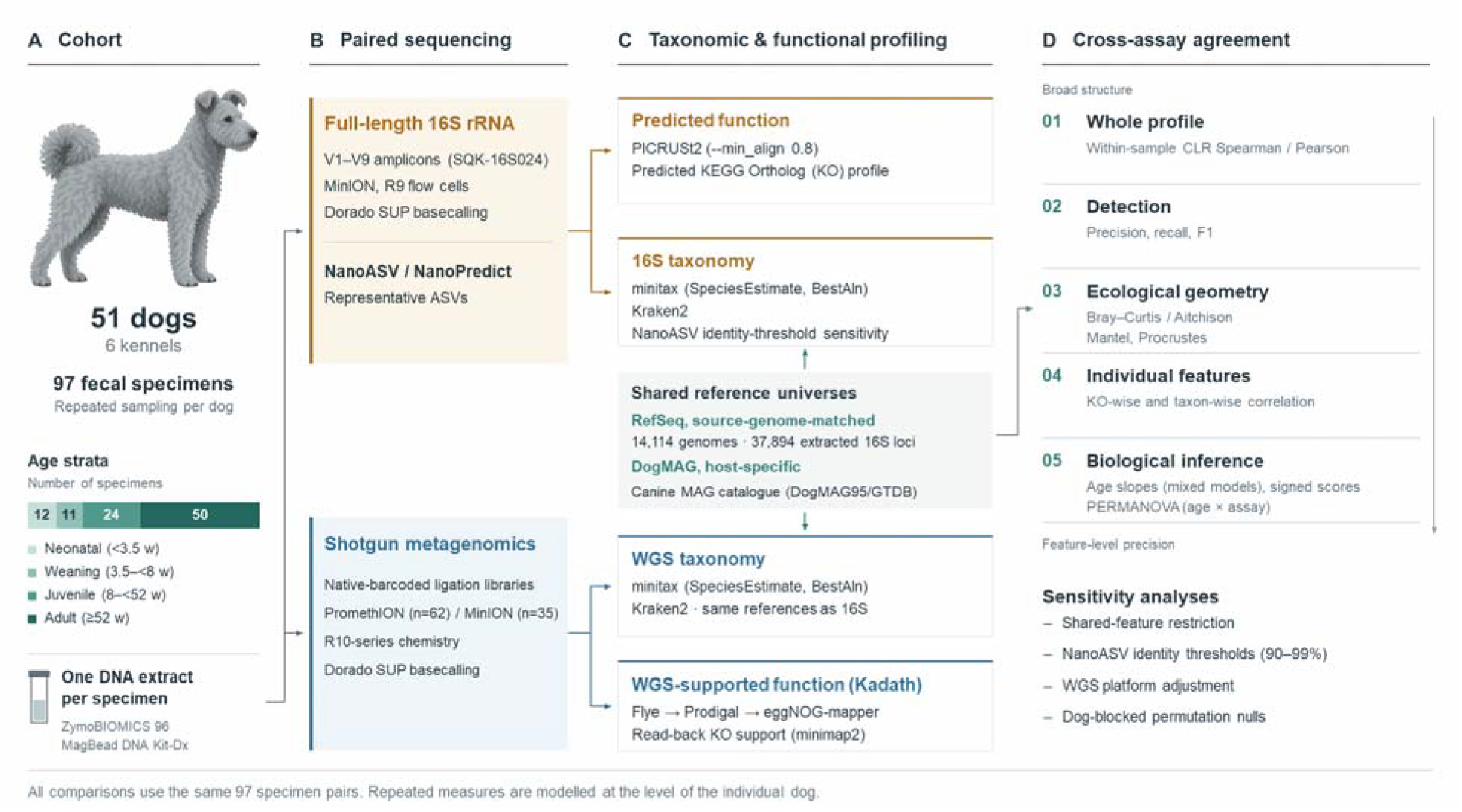
Study design and analytical workflow. Ninety-seven matched canine fecal specimens from 51 verified dogs were profiled by full-length ONT 16S and ONT WGS. Functional analyses compared NanoASV/NanoPredict-PICRUSt2 predictions with Kadath WGS-supported KO profiles at whole-profile, individual-feature and biological-inference scales. Taxonomic analyses compared source-genome-matched RefSeq profiles, dog-aware ecological structure, DogMAG reference harmonization, multivariate developmental inference and NanoASV sequence-matching sensitivity.

**Figure 2:**
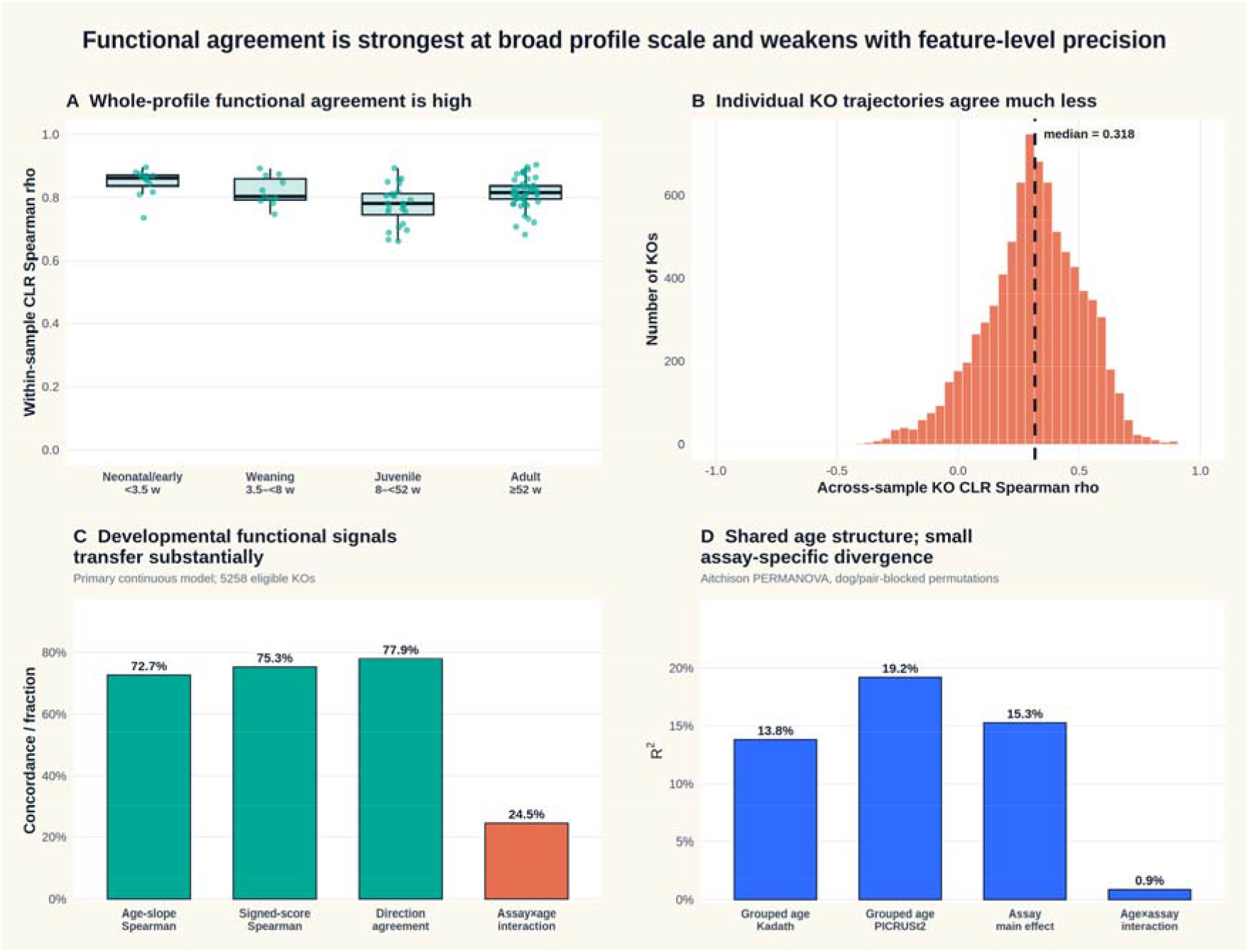
Functional concordance across analytical scales in the 97-pair cohort. Whole-profile CLR agreement is high and between-sample functional structure is preserved, whereas feature-wise KO abundance transfer is substantially weaker. Continuous age-associated KO effects show considerable directional and slope concordance but also show frequent assay-by-age interactions, with closer agreement in broad biological patterns than in feature-level effect magnitudes.

**Figure 3:**
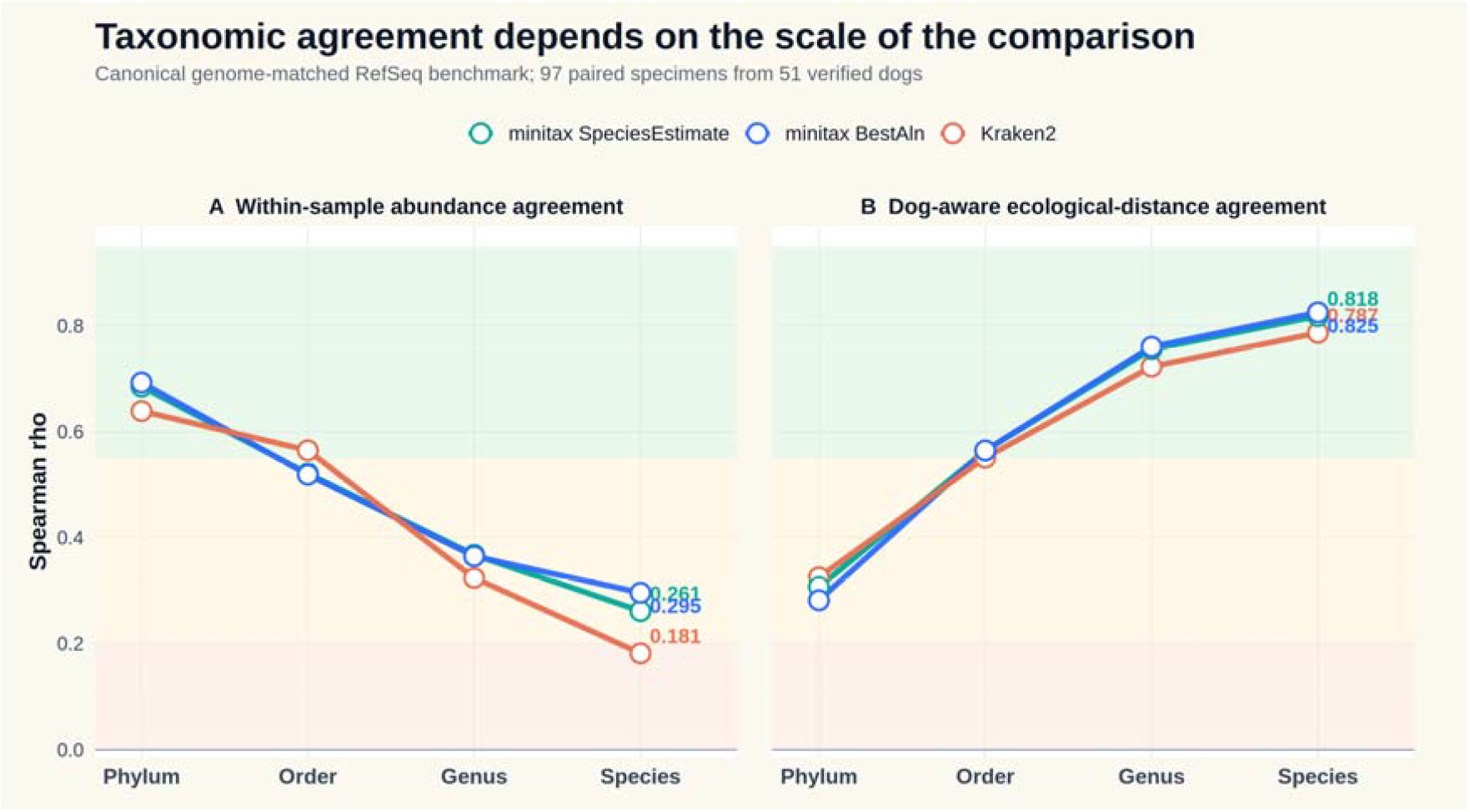
Taxonomic concordance depends strongly on the scale of comparison. In the source-genome-matched RefSeq benchmark, exact within-sample fine-rank abundance agreement is modest, whereas dog-aware between-sample ecological distance concordance remains high. Agreement in relationships among specimens therefore exceeds agreement in genus- or species-level abundances.

**Figure 4:**
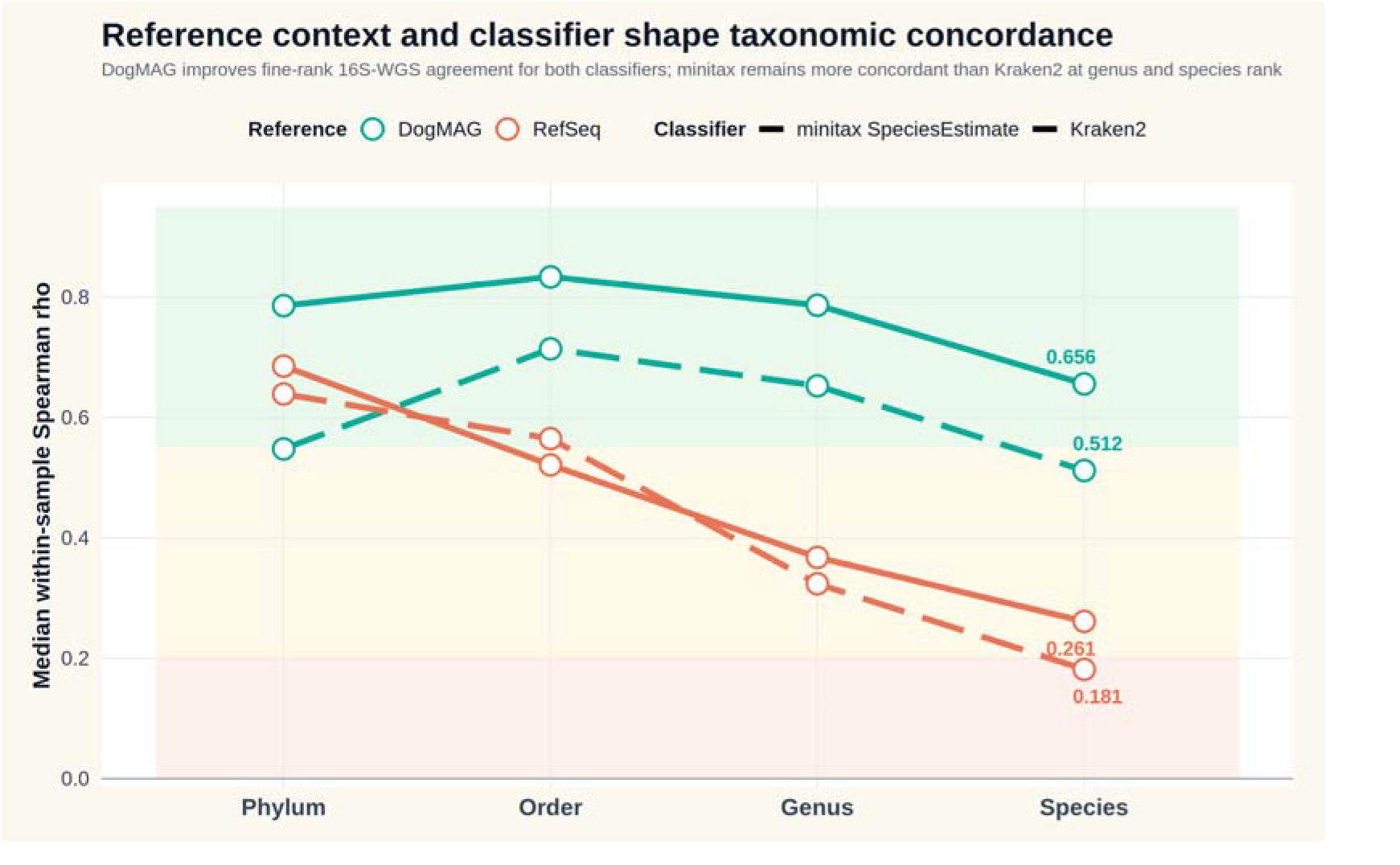
Reference context and classifier jointly shape taxonomic concordance. Median within-sample Spearman correlations are shown across taxonomic ranks for minitax SpeciesEstimate and Kraken2 under the source-genome-matched RefSeq benchmark and the host-specific DogMAG reference. DogMAG increases genus/species concordance for both classifiers, consistent with an effect of reference representation, while minitax remains more concordant than Kraken2 under both matched reference contexts, indicating an additional classifier effect.

**Figure 5:**
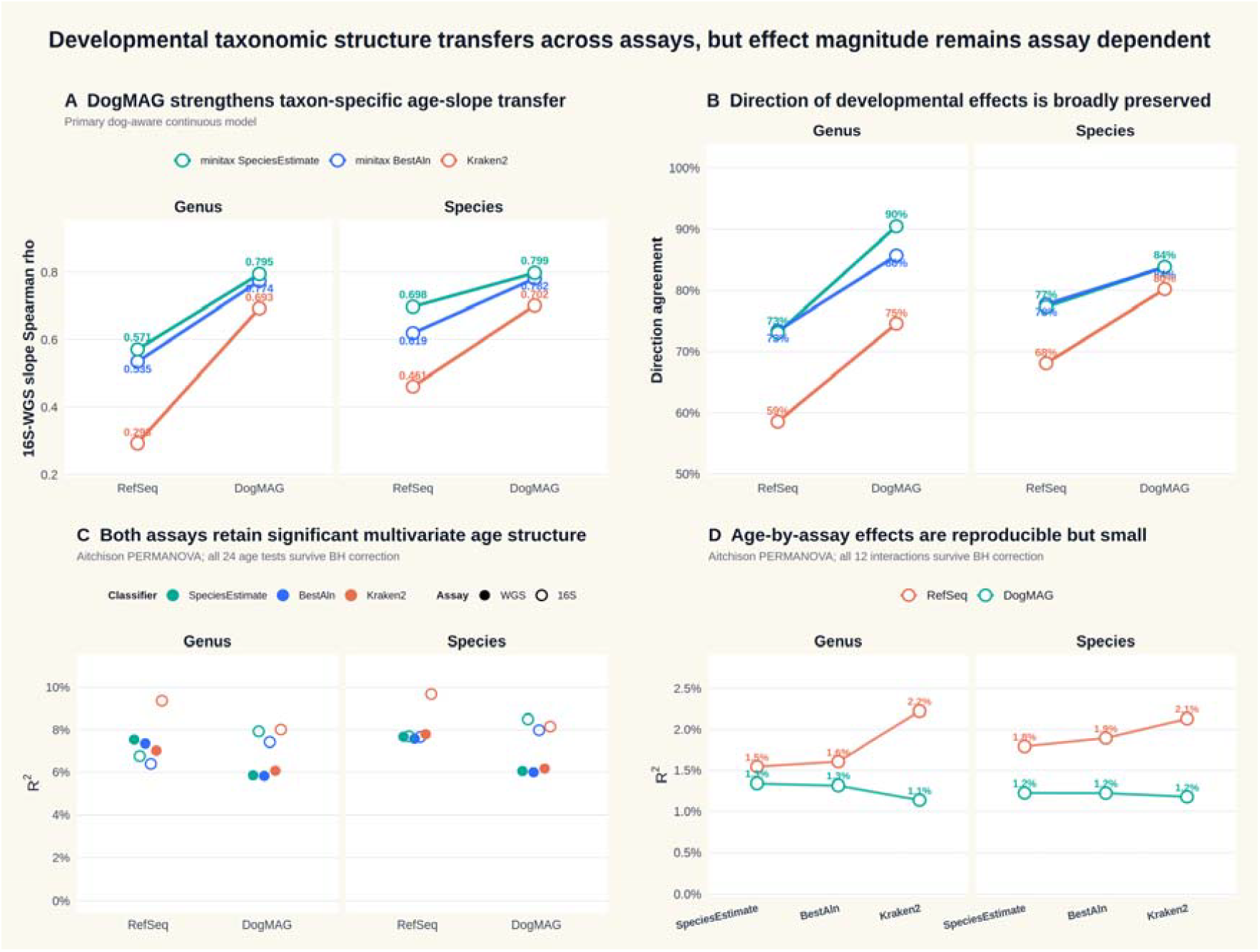
Transfer of age-associated taxonomic structure across assays. Continuous dog-aware taxon slopes show stronger 16S–WGS concordance under DogMAG than RefSeq for every classifier, while multivariate PERMANOVA shows significant developmental structure in both assays. Paired age-by-assay effects are consistently significant but small relative to the main assay difference, indicating broad transfer of developmental organization without equivalence of taxon-specific effect magnitude.

**Figure 6:**
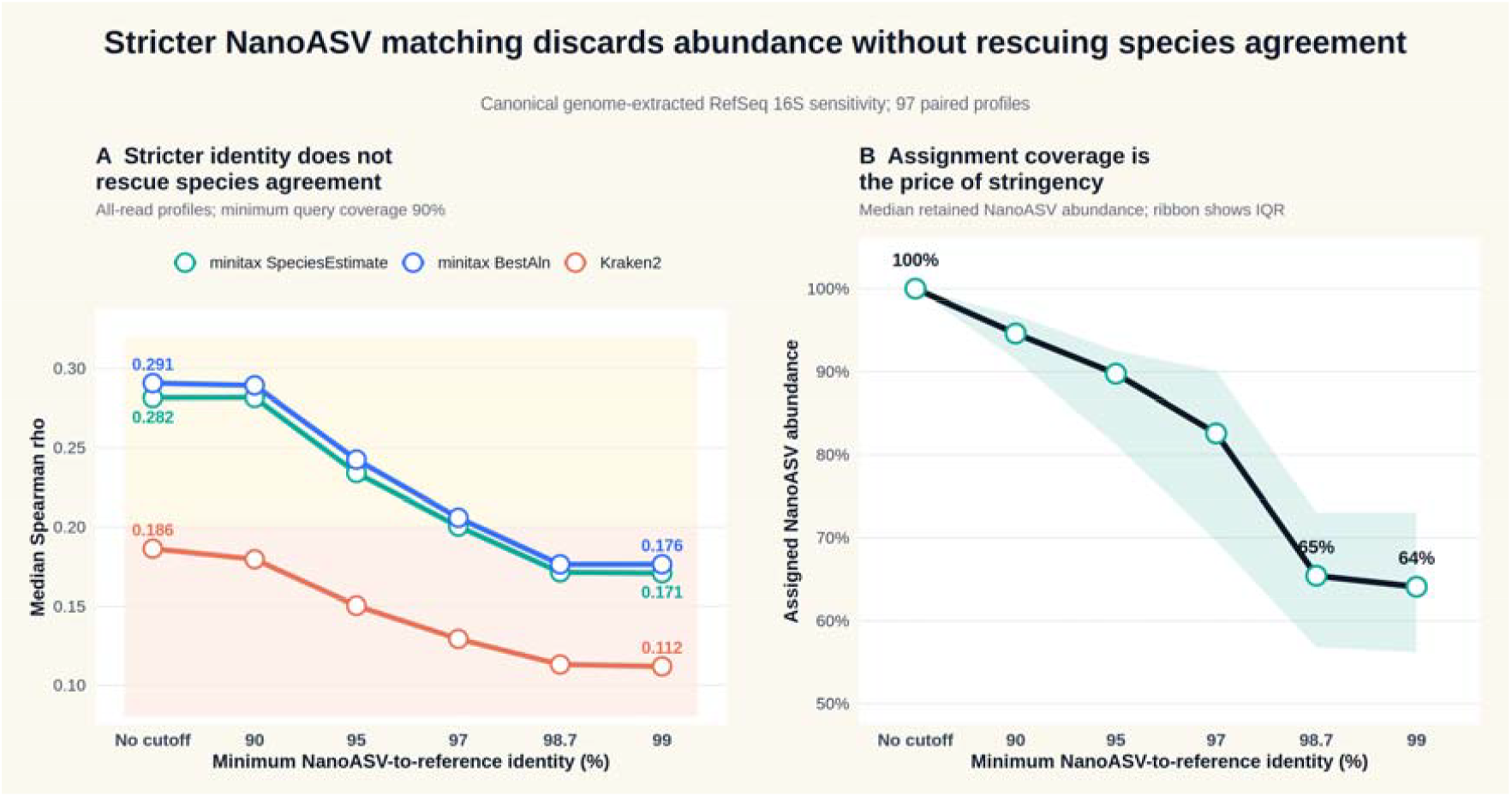
Stricter NanoASV-to-reference sequence matching removes read abundance without materially improving species-level agreement across all 97 matched profiles. Increasing sequence-identity stringency progressively reduces retained assigned abundance, while species-level Spearman concordance remains unchanged at mild filtering and then declines. Hard identity cutoffs did not reduce the fine-rank discrepancy between assays.

**Figure 7:**
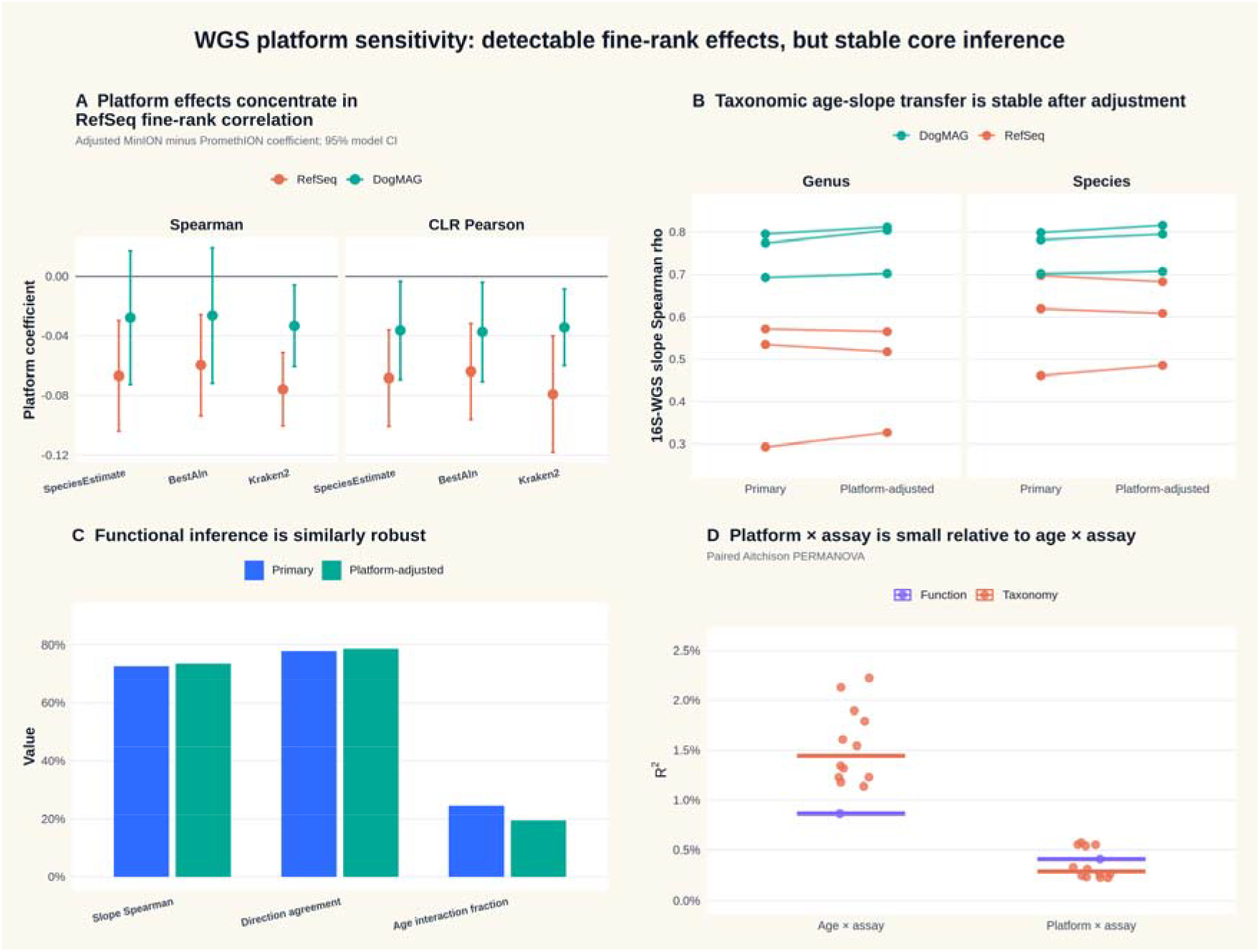
WGS-platform sensitivity summary. Platform-associated effects are strongest for fine-rank RefSeq abundance correlations and are markedly attenuated under DogMAG. Platform adjustment leaves continuous age-signal transfer nearly unchanged, while paired assay-by-platform PERMANOVA interactions remain small and non-significant compared with significant age-by-assay interactions. Because platform was not randomized, these estimates are technical robustness checks rather than causal instrument effects.

### ONT shotgun metagenomic sequencing

WGS libraries were prepared from the same DNA extracts that were used for full-length 16S sequencing (ZymoBIOMICS 96 MagBead DNA Kit-Dx; see below), so both assays were applied to identical nucleic-acid material from each specimen. Native-barcoded ONT ligation libraries were prepared from these extracts. The original WGS branch was sequenced on PromethION using R10-series chemistry; the added branch used MinION while retaining the same general extraction, library-preparation and basecalling framework. Basecalling was performed with Dorado v0.8.3 in super-accurate mode followed by barcode demultiplexing. Library identity, sequencing branch and specimen of origin are recorded in the specimen-pairing metadata.

### ONT full-length 16S sequencing

The matched full-length 16S data derive from the longitudinal canine ONT 16S study described by Asaduzzaman et al. (Asaduzzaman et al. 2026). Fecal material was collected using DNA/RNA Shield Fecal Collection Tubes and stored at −80 degrees C. DNA was extracted with the ZymoBIOMICS 96 MagBead DNA Kit-Dx (D4308-E), quantified with a Qubit 4 fluorometer and assessed with an Agilent 4150 TapeStation. Full-length V1–V9 16S libraries were prepared using the ONT SQK-16S024 workflow with AMPure XP cleanup and sequenced on MinION R9 flow cells. Reads were basecalled with Dorado v0.8.0 in super-accurate mode with a minimum Q score of 10 (Asaduzzaman et al. 2026). Only libraries from specimens with a matched WGS library were retained here.

### NanoASV processing and PICRUSt2 functional prediction

Full-length 16S reads were processed with the NanoASV/NanoPredict workflow to generate representative ASV sequences and sample-specific ASV read-count tables. Sample identifiers were reconciled to the specimen metadata before downstream analysis. NanoASV abundance profiles were supplied to PICRUSt2 (Douglas et al. 2020), using the placement setting `--min_align 0.8`. Predicted KO abundances were taken from the unstratified `KO_metagenome_out/pred_metagenome_unstrat.tsv.gz` output and standardized to KEGG Ortholog identifiers.

PICRUSt2 values are interpreted as phylogenetically predicted gene-family abundance rather than direct observation. They depend on reference phylogeny, inferred genome content and marker-gene copy-number modeling and cannot directly recover sample-specific gene gain or loss, mobile elements or strain-level accessory functions.

### WGS functional profiling with Kadath

Kadath provided the specimen-level WGS functional profiles. Each WGS library was assembled independently with Flye in metagenome mode using ONT high-quality reads (`--meta --nano-hq`) (Kolmogorov et al. 2019). Coding sequences were predicted with Prodigal in metagenomic mode (`-p meta`) (Hyatt et al. 2010). Predicted proteins were annotated with eggNOG-mapper using DIAMOND searches (`--itype proteins -m diamond`), and the `KEGG_ko` field was retained as the common functional vocabulary (Cantalapiedra et al. 2021; Kanehisa et al. 2021).

The original WGS reads for each specimen were mapped back to that specimen’s assembly with minimap2 using the `map-ont` preset (Li 2018). Mapped primary alignments were retained after excluding unmapped, secondary and supplementary records; no minimum mapping-quality filter was applied (MAPQ ≥ 0). Alignments were coordinate sorted and indexed, and read overlaps with predicted genes were counted with sorted `bedtools coverage-counts`. Gene-level read support was joined to eggNOG-mapper annotations and aggregated by KO. When a gene carried multiple KO annotations, the primary analysis assigned its full supported count to each annotation; fractional multi-KO assignment and alternative length-normalized abundance definitions were retained as sensitivity analyses (Tamames and Puente-Sánchez 2019; Franzosa et al. 2018).

Kadath estimates depend on assembly, gene prediction, annotation and read support. A sequence must assemble, be predicted as a gene, receive an orthology annotation and receive read-back support to contribute to the final WGS KO matrix. Failure at one of these stages is not interpreted as proof that the function is biologically absent.

### Functional concordance and inference-transfer analysis

PICRUSt2 and Kadath KO tables were paired by specimen identifier and KO for all 97 matched specimens. The merged union contained 9,761 KOs; 8,225 passed the primary prevalence filter requiring nonzero relative abundance in at least two paired specimens in either method. Within each method, KO counts were converted to relative abundance. CLR transformations used a relative-abundance pseudocount of 10^−8^ (Aitchison 1982).

Functional agreement was evaluated at progressively stricter analytical scales. Whole-profile agreement was measured within each specimen using Pearson and Spearman correlations of CLR-transformed KO vectors. Between-sample functional structure was compared with Euclidean distances in CLR space, Spearman Mantel correlation (Mantel 1967) and Procrustes concordance with vegan::protest; permutation-based tests used 999 permutations. Individual KOs were then followed across specimens to quantify feature-wise abundance transfer. Detection overlap used relative abundance >10^−4^ as the primary presence threshold, with zero as a sensitivity.

Age-associated functional inference was evaluated in both continuous and grouped forms. The primary continuous analysis retained KOs detected in at least ten specimens in both assays and fitted standardized log1p(age_weeks); standardized raw age in weeks was retained as a sensitivity. For each assay separately, KO CLR abundance was modeled with age and kennel as fixed effects and dog as a random intercept. Joint paired models included an assay-by-age interaction with specimen pair nested within dog, testing whether PICRUSt2 and Kadath estimated different age slopes for the same KO. Transfer was summarized by Spearman/Pearson correlation of age slopes, signed age scores sign(*β*)× −log_10_ (p), direction agreement and 999 KO-label permutations. The grouped analysis used the four developmental strata and the signed-score framework adapted from Sun et al. (Sun, Jones, and Fodor 2020), with neonatal/early specimens as the prespecified reference and 999 age-label and KO-label permutations.

Multivariate functional inference was evaluated with PERMANOVA. Aitchison distance was primary and Bray–Curtis was a sensitivity. Within each assay, continuous age and grouped age effects were tested with permutations blocked by verified dog identity; kennel and WGS acquisition platform were included as nuisance terms for the WGS profiles, and the paired WGS platform label was carried as a nuisance stratum for the matched 16S profile. Paired assay effects and assay-by-age interactions were tested after stacking the 16S and WGS profiles, with permutations restricted within biological specimen pair. These analyses distinguish preservation of broad multivariate biological structure from agreement in individual KO effects.

### Source-genome-matched RefSeq taxonomy

Taxonomic concordance was evaluated under a controlled reference design in which both assays were constrained to the same underlying source-genome universe. The reference contained 14,114 bacterial and archaeal RefSeq reference/representative assemblies (13,561 bacterial and 553 archaeal; complete genome, chromosome or scaffold level). Full-length 16S loci were extracted from these same source assemblies, yielding 37,894 retained 16S sequences from 13,187 genomes; 927 source genomes had no retained 16S locus.

WGS and 16S profiles were generated against this harmonized source-genome universe with minitax (Gulyás et al. 2024), using both `SpeciesEstimate` and `BestAln` abundance definitions, and with Kraken2 as a complementary classifier (Wood, Lu, and Langmead 2019). Concordance was summarized for both all-read profiles, which retained unclassified abundance, and classified-only profiles, in which classified taxa were renormalized. Primary within-sample summaries included Spearman correlation, CLR Pearson correlation, Bray– Curtis dissimilarity and detection F1 across taxonomic ranks. To test classifier effects directly in the RefSeq benchmark, specimen-level minitax-minus-Kraken2 differences were reduced to one mean contrast per verified dog and evaluated by dog-level sign-flip permutations with dog bootstrap confidence intervals. The same repeated-dog principle was used for the formal DogMAG classifier comparison; effect orientation was defined so that positive DogMAG advantages consistently indicated stronger minitax concordance.

### Taxonomic inference-transfer analysis

To distinguish taxonomic profile similarity from reproducibility of biological conclusions, the signed inference-transfer framework used for KOs was generalized to genus- and species-level taxonomic abundance tables for all three classifiers/estimators. Categorical analyses used the developmental strata defined in weeks: neonatal/early (<3.5 weeks; n=12), weaning (3.5–<8 weeks; n=11), post-weaning/juvenile (8–<52 weeks; n=24) and adult (>=52 weeks; n=50), with the neonatal/early group as the prespecified reference. Taxa were required to be detected in at least five specimens in both assays. For each eligible taxon and contrast, a Wilcoxon test was run independently in 16S and WGS and the direction of the abundance difference was combined with statistical evidence as *S* = sign(*Δ*) × −log_10_ (p). Matched 16S and WGS score vectors were compared by Spearman and Pearson correlation plus direction agreement. Age-label permutations generated the primary null, while taxon-label permutations tested whether concordance depended on matching the same biological taxon; 999 permutations were used for each null.

The categorical analysis was run under the source-genome-matched RefSeq benchmark for minitax `SpeciesEstimate`, minitax `BestAln` and Kraken2 and repeated in DogMAG for both minitax estimators and Kraken2. For direct RefSeq-versus-DogMAG comparison, the primary matrix uses aligned nonzero prevalence eligibility. DogMAG Kraken2 additionally retains a >=0.01% relative-abundance eligibility sensitivity because confidence-zero classification against the compact DogMAG panel produces diffuse low-abundance assignments.

Continuous age was treated as the primary inferential analysis because it retains the developmental gradient and can model repeated sampling explicitly. Classified taxa were renormalized within specimen and rank and transformed to CLR values using a relative-abundance pseudocount of 10^−6^. Taxa were retained when detected in at least ten specimens in both assays. Standardized log(1 + age in weeks) was the primary age covariate; standardized raw age in weeks was a sensitivity. For each assay separately, taxon-level CLR abundance was fitted as `CLR ~ age + kennel` with a random intercept for dog. Joint models used `CLR ~ assay * age + kennel` with sample pair nested within dog as the random structure; the assay-by-age interaction tested whether age slopes differed between 16S and WGS. Cross-assay transfer was summarized from the corresponding age slopes, signed age scores sign(*β*) × −log_10_ (*p*), direction agreement and taxon-label permutation tests. These models estimate concordance of age-associated structure and assay-specific slope differences; they do not establish causal developmental effects.

### Dog-aware taxonomic structure and metadata-associated effects

Repeated sampling was addressed using verified dog identity. Cross-assay Bray–Curtis distance concordance was tested with permutations restricted within dog, using all 97 pairs from 51 dogs. The continuous taxonomic age models described above included dog-level random effects and kennel adjustment, with a joint assay-by-age model used to identify taxa whose estimated developmental slopes differed between assays.

Multivariate taxonomic inference used prevalence-filtered classified-only genus and species profiles. Taxa were retained when relative abundance exceeded 10^−4^ in at least 10% of specimens in either assay, and retained profiles were renormalized. Aitchison distance was primary and Bray–Curtis was used as a sensitivity. Within-assay continuous and grouped age effects were tested with dog-blocked permutations. Paired assay effects and assay-by-age interactions were tested on stacked WGS/16S profiles with permutations restricted within specimen pair. PERMANOVA was repeated for RefSeq and DogMAG, minitax SpeciesEstimate, minitax BestAln and Kraken2 at genus and species rank. These multivariate tests complement the feature-level mixed models by asking whether broad developmental community structure transfers even when individual taxa show different effect magnitudes.

### DogMAG host-specific reference sensitivity

To test whether generic reference coverage contributed to fine-rank disagreement, both 16S and WGS were additionally analyzed in a host-specific DogMAG reference context (Kakuk et al. 2026b). The minitax analysis used `SpeciesEstimate` and `BestAln` profiles for the full 97-pair set. Shared-taxa analyses, WGS genome-size correction and MAG-derived 16S copy-number correction were evaluated as sensitivities.

A matched Kraken2 analysis classified the same 97 WGS libraries and their corresponding 97 full-length 16S libraries against the identical DogMAG95/GTDB database at confidence 0. `Specimen pairing was taken from explicit run-level sample manifests rather than inferred from file names.` Kraken2 report counts were propagated through the report hierarchy to obtain comparable rank-level profiles; reads not resolved to a requested rank contributed to the unclassified component at that rank. The same primary concordance framework was then applied to minitax and Kraken2: per-sample relative-abundance and CLR correlations, detection precision/recall/F1, Bray–Curtis and Aitchison distance concordance, feature-space sensitivities, PCoA and taxon-level age-signal transfer. The primary detection threshold was relative abundance 10^−4^. Restriction to taxa observed in both assays was used to quantify how much disagreement arose from assay-specific feature detection rather than abundance differences among shared taxa.

This matched design provides two complementary tests. Comparing RefSeq with DogMAG within a classifier estimates the effect of reference context, whereas comparing minitax with Kraken2 within DogMAG estimates the effect of classifier behavior while holding the host-specific reference universe and specimen set constant.

### Read-count-weighted NanoASV sequence-matching sensitivity

A separate sequence-stringency experiment was run on all 97 matched profiles to test whether poor species-level concordance could be explained mainly by permissive matching of NanoASV representative sequences. Each matched pair was linked to its NanoASV per-barcode abundance file, preserving the original sample-specific read count associated with each representative sequence. NanoASV representatives were aligned with minimap2 against the genome-extracted 16S reference. Identity thresholds of 90%, 95%, 97%, 98.7% and 99% were tested at minimum query coverages of 90% and 95%; unrestricted best-hit assignment provided the baseline. The same threshold definitions, presence threshold (relative abundance 1e-4), CLR pseudocount (1e-6) and WGS comparators (minitax `SpeciesEstimate`, minitax `BestAln` and Kraken2) were retained from the historical 62-pair sensitivity so that cohort expansion could be evaluated without changing the analysis definition.

For each threshold and taxonomic rank, the reads carried by ASVs that passed the sequence criterion contributed to the matched reference taxon. Reads associated with ASVs failing the threshold or lacking a rank-level label were assigned to `Unclassified`. Taxonomic profiles were reconstructed by summing the original ASV read counts within each assigned taxon. All-read profiles therefore preserved failed-assignment abundance in the `Unclassified` compartment, whereas classified-only profiles removed `Unclassified` and renormalized the remaining taxa. This analysis tests whether stricter sequence matching produces a smaller but quantitatively more WGS-concordant 16S profile.

### WGS platform sensitivity analysis

The 97 WGS profiles comprised 62 PromethION and 35 MinION libraries. Platform was not randomized: the MinION subset was enriched for younger specimens, although both platforms were represented in all six kennels. Platform-associated estimates are therefore treated as observational technical sensitivities rather than causal instrument effects.

Functional sensitivity analyses repeated the continuous KO models with WGS platform added to assay-specific models and an assay-by-platform term added to joint paired models. Grouped age-label permutations were additionally stratified within WGS platform. The functional PERMANOVA reported the paired assay-by-platform interaction separately from the assay-by-age interaction.

Taxonomic sensitivity analyses used the same strategy. Specimen-level concordance metrics were modeled as metric ~ platform + log1p(age) + kennel + (1|dog). Grouped taxonomic age-label permutations were restricted within platform strata. Continuous taxon models added WGS platform to assay-specific models and assay-by-platform to paired joint models. Taxonomic PERMANOVA likewise separated assay-by-platform from assay-by-age interactions. Non-platform-adjusted taxonomy results are reported as the primary estimates; the platform-adjusted analyses test robustness to the mixed WGS acquisition history.

## Results

### Ninety-seven matched specimens were included in both functional and taxonomic analyses

The analytical set comprised 97 matched ONT 16S–WGS specimens from 51 verified dogs distributed across six kennels. The same 97 specimen pairs were used for the functional and taxonomic comparisons. Developmental strata contained 12 neonatal/early, 11 weaning, 24 post-weaning/juvenile and 50 adult specimens.

### Functional profiles agreed strongly at the whole-community level

The PICRUSt2–Kadath comparison contained 9,761 KOs before prevalence filtering and 8,225 KOs after the primary filter. Whole-profile concordance was consistently high. Median within-sample CLR Spearman correlation was 0.860 in neonatal/early specimens, 0.804 in weaning, 0.781 in post-weaning/juvenile and 0.816 in adults; the corresponding CLR Pearson medians ranged from 0.774 to 0.867.

Between-sample functional structure was also preserved. The Mantel Spearman correlation between PICRUSt2 and Kadath distance matrices was 0.543 and the Procrustes correlation was 0.693 (both p=0.001, 999 permutations). At the primary 0.01% presence threshold, median detection F1 ranged from 0.772 to 0.831 across developmental strata. These results indicate agreement in both broad functional composition and relationships among specimens.

### Individual KO abundances were less concordant than whole profiles

Feature-wise agreement was markedly weaker than sample-wise profile agreement. Across 8,225 prevalence-filtered KOs, the median KO-wise CLR Spearman correlation was 0.318 (IQR 0.195–0.449), although 92.9% of KO correlations were positive. Whole-profile similarity thus reflects broad functional composition and relationships among samples; individual KO trajectories can still differ between assays.

### Age-associated functional effects were concordant, with differences in magnitude

Among 5,258 KOs eligible for the primary nonzero continuous-age analysis, PICRUSt2 and Kadath age slopes were substantially concordant (Spearman=0.727; Pearson=0.767). Signed age-score concordance was 0.753 by Spearman and direction agreement was 77.9%. Age-associated evidence was individually significant at FDR<0.05 for 2,496 Kadath KOs and 3,843 PICRUSt2 KOs; 2,112 were significant in both methods and 2,052 of those shared the same direction. All primary KO-label permutation tests gave q=0.001, showing that the transfer depended on matching the same KO identities rather than arbitrary feature pairing.

The paired joint model identified significant assay-by-age interactions for 1,290/5,258 eligible KOs (24.5%). At the stricter 0.01% eligibility threshold, slope concordance remained positive but fell to 0.606 across 2,493 KOs, with 617 significant assay-by-age interactions. Developmental functional signals were concordant overall, although feature-level effect magnitudes differed.

In the multivariate analysis, the paired assay effect was strong (Aitchison R2=0.153, p=0.001; Bray–Curtis R2=0.278, p=0.001), confirming a systematic difference between the functional profiles produced by the two assays. Grouped age structure was significant within both methods (Aitchison R2=0.138 for Kadath and 0.192 for PICRUSt2; p=0.003 and 0.004). Continuous age explained a smaller fraction of multivariate variation and was metric-dependent (Aitchison R2=0.044, p=0.049 for Kadath and R2=0.096, p=0.058 for PICRUSt2). The paired age-by-assay interaction was significant but small (Aitchison R2=0.0086, p=0.025; Bray–Curtis R2=0.0140, p=0.014), indicating broadly similar developmental trajectories with some assay-specific differences.

### RefSeq taxonomy preserved ecological structure better than exact fine-rank abundance

Under the source-genome-matched RefSeq benchmark, exact within-sample abundance agreement declined strongly toward fine ranks. Classified-only median species-level Spearman correlation was 0.261 for minitax SpeciesEstimate, 0.295 for BestAln and 0.181 for Kraken2. Genus-level values were higher but still moderate.

Agreement was higher for between-sample ecological distances. Dog-aware species-level Bray–Curtis distance concordance was 0.818 for SpeciesEstimate, 0.825 for BestAln and 0.787 for Kraken2 (all permutation q approximately 0.001). The assays agreed more closely on relationships among specimens than on species proportions within each specimen.

### DogMAG improved taxonomic concordance for both classifiers; minitax remained more concordant than Kraken2

Using DogMAG in place of RefSeq increased species-level median within-sample Spearman correlation from 0.261 to 0.656 for minitax SpeciesEstimate and from 0.181 to 0.512 for Kraken2. At genus rank, the corresponding changes were 0.367 to 0.786 and 0.324 to 0.652. The improvement with the host-specific reference was observed with both classifiers.

Classifier differences were also evident within each reference set. With RefSeq, SpeciesEstimate outperformed Kraken2 for all eight prespecified genus/species paired endpoints spanning Spearman correlation, CLR Pearson correlation, detection F1 and Bray– Curtis distance after BH correction. With DogMAG, all 10 prespecified SpeciesEstimate-versus-Kraken2 primary endpoints across abundance agreement, detection performance and within-specimen distance were significant after BH correction, and the BestAln sensitivity showed the same direction. Minitax therefore showed stronger cross-assay concordance across the tested endpoints and both references. This comparison does not establish universal classifier accuracy, because it lacks an external taxonomic ground truth.

Restricting DogMAG Kraken2 to taxa observed by both assays increased species-level Spearman from 0.512 to 0.666 and detection F1 from 0.726 to 0.892. Differential feature detection therefore accounts for a substantial component of the fine-rank gap, although abundance disagreement remains among shared taxa.

### Age-associated taxonomic patterns were concordant despite differences in effect magnitude

The primary dog-aware continuous models showed that DogMAG strengthened transfer of taxon-specific age effects for every classifier/estimator. For minitax SpeciesEstimate, genus/species slope Spearman increased from 0.571/0.698 under RefSeq to 0.795/0.799 under DogMAG. BestAln increased from 0.535/0.619 to 0.774/0.782, and Kraken2 from 0.293/0.461 to 0.693/0.702. All primary taxon-label permutation tests remained significant at q=0.001. Minitax therefore retained stronger age-slope transfer than Kraken2 under both reference contexts.

The multivariate analysis also detected developmental structure. With Aitchison distance, continuous age was significant after multiple-testing correction in all 24 assay/reference/classifier/rank combinations and explained approximately 5.8–9.7% of community variation. The four-group developmental factor was significant in all 24 combinations and explained approximately 14–24%. The paired assay effect was significant in all 12 reference/classifier/rank combinations (q=0.001), and age-by-assay interactions were significant in all 12 while accounting for only about 1.1–2.2% of Aitchison variation. The assays captured similar developmental structure, with smaller differences in their age-associated trajectories.

In the feature-level mixed models, significant assay-by-age interactions remained common even when global slope rankings were strongly concordant; for DogMAG SpeciesEstimate, 33/63 eligible genus taxa and 63/99 species taxa showed significant interactions in the primary model.

The grouped signed-score analysis provided an independent categorical sensitivity with explicit permutation nulls (Supplementary Figure S1). Under RefSeq, all 18 classifier × rank × developmental-contrast tests exceeded the age-label permutation null after BH correction, and every taxon-label permutation test was significant at q=0.001. Under DogMAG minitax, 8/12 primary grouped tests were age-label significant: the juvenile-versus-neonatal and adult-versus-neonatal contrasts were significant for both SpeciesEstimate and BestAln at genus and species rank, whereas the earliest weaning-versus-neonatal contrast was not significant after BH correction. Under primary nonzero DogMAG Kraken2, observed grouped concordance remained high but none of the six age-label tests survived BH correction; nevertheless, all six taxon-label permutation tests remained significant at q=0.001. Taxon-label significance shows that concordance depends on matching the correct biological taxa, whereas age-label significance asks whether the observed grouped developmental concordance exceeds what is expected from shuffled age assignments.

### Stricter 16S sequence matching did not improve species-level agreement

The 97-pair NanoASV sequence-stringency sensitivity tested whether residual species disagreement was driven mainly by weak ASV-to-reference matches. With 90% minimum query coverage, the median fraction of NanoASV abundance remaining assigned was 94.6% at 90% identity, 89.8% at 95%, 82.6% at 97%, 65.4% at 98.7% and 64.1% at 99%.

Species-level agreement did not improve as matching became more stringent. Against WGS SpeciesEstimate, median Spearman was 0.282 with unrestricted best-hit assignment, remained essentially unchanged at 90% identity, and then declined to 0.234, 0.200, 0.171 and 0.171 at 95%, 97%, 98.7% and 99%. BestAln and Kraken2 showed the same qualitative decline. Hard identity filtering therefore removes substantial marker-gene abundance without moving the retained profile closer to WGS.

### WGS platform adjustment had little effect on cross-assay comparisons

WGS platform was associated with some abundance-correlation metrics. After adjustment for age, kennel and repeated dog sampling, MinION-associated WGS profiles showed lower RefSeq fine-rank abundance correlations with 16S. For example, the MinION-versus-PromethION coefficient for species-level Spearman was −0.067 for RefSeq SpeciesEstimate (q=0.0054), −0.060 for BestAln (q=0.0063) and −0.076 for Kraken2 (q=1.4e-5). CLR Pearson showed the same pattern, whereas F1 and Bray–Curtis generally did not. Under DogMAG these platform associations were markedly attenuated; only Kraken2 species-level CLR Pearson remained BH significant (q=0.045).

Platform adjustment did not materially change age-signal transfer. RefSeq SpeciesEstimate genus/species slope concordance changed from 0.571/0.698 to 0.565/0.683, whereas DogMAG SpeciesEstimate changed from 0.795/0.799 to 0.812/0.816. DogMAG Kraken2 changed from 0.693/0.702 to 0.702/0.707. The proportion of taxa with significant assay-by-age interactions generally decreased modestly after adjustment, indicating that acquisition branch explains part, but not most, of the assay-specific age-effect differences.

No taxonomic assay-by-platform interaction was significant after multiple-testing correction in any of the 12 Aitchison comparisons (R2 approximately 0.22–0.57%; all q=0.441), whereas every age-by-assay interaction remained significant. Functional results were similarly stable: the primary KO slope Spearman changed from 0.727 to 0.736 after platform adjustment, and direction agreement from 77.9% to 78.7%. The functional assay-by-platform interaction was non-significant by both Aitchison (R2=0.0041, p=0.279) and Bray–Curtis (R2=0.0017, p=0.531). Platform therefore contributes to some fine-rank RefSeq abundance discrepancies but does not explain the main reference, classifier, assay or developmental-transfer effects.

## Discussion

Across the 97 matched specimens, full-length ONT 16S and ONT WGS agreed closely on broad functional composition, relationships among samples and developmental patterns. Agreement was lower for fine-rank abundances and individual-feature effect magnitudes. This dependence on analytical scale helps define the uses for which 16S can serve as an alternative to WGS.

Whole-sample KO profiles and between-sample functional relationships were concordant, while individual KO abundance trajectories were less reproducible. Age slopes and signed age scores also agreed substantially across more than five thousand eligible KOs, with approximately 78% directional agreement. Roughly one quarter of eligible KOs retained significant assay-by-age interactions, and the paired multivariate age-by-assay effect was significant despite explaining only about 1% of functional variation. PICRUSt2 therefore captures much of the direction and ordering of developmental functional change while remaining unreliable as a quantitative substitute for WGS-supported KO effect sizes. This scale dependence is consistent with the distinction between phylogenetically expected gene content and directly supported sample-specific sequence content (Douglas et al. 2020; Sun, Jones, and Fodor 2020; Matchado et al. 2024).

For taxonomy, the difference between levels of analysis was larger. Under the source-genome-matched RefSeq benchmark, exact species-level abundance agreement was modest, yet between-sample ecological structure remained strong. In the PERMANOVA analysis, continuous age explained a reproducible fraction of Aitchison community variation in every assay/reference/classifier/rank combination, while the paired age-by-assay interactions were consistently significant but much smaller than the main assay effect. Both assays therefore detect the same broad developmental restructuring of the canine gut microbiome, but they do not trace identical multivariate trajectories or assign identical taxon-specific effect magnitudes.

Reference representation contributed to these differences. DogMAG increased fine-rank abundance concordance for both minitax and Kraken2 and substantially strengthened transfer of continuous taxon-specific age slopes. Because both classifiers improve when moved into the same host-specific reference set, the gain is not specific to a minitax estimator. DogMAG should nevertheless be interpreted as host-specific reference harmonization rather than independent external validation: it derives from the same broader canine sampling effort and intentionally restricts the comparison to a feature space better represented in dogs.

Classifier choice affected agreement independently of reference choice. Minitax consistently produced stronger 16S–WGS concordance than Kraken2 under both RefSeq and DogMAG, and the formal dog-aware paired tests supported that ordering across every prespecified primary abundance, detection and within-specimen distance endpoint tested in the two matched reference settings. Minitax also showed stronger genus-and species-level age-slope transfer than Kraken2 under both reference contexts. Because the comparison holds specimen set and reference set constant, these differences suggest that the handling of classification evidence and abundance estimation affect cross-assay agreement. This advantage is specific to concordance in the present benchmark. WGS served as an operational comparator, so the results do not establish universal taxonomic accuracy.

Differences in taxon detection also contributed. Restricting DogMAG comparisons to taxa detected by both assays substantially improves species-level correlation and detection F1, demonstrating that disagreement often arises before abundance estimation, at the level of which taxa are represented at all. Marker-gene resolution is constrained by locus conservation, amplification and rRNA operon biology, whereas WGS draws information from across genomes. Abundance discrepancies remained among shared taxa, indicating additional sources of disagreement.

The NanoASV analysis tested whether permissive low-identity ASV matches contributed to poor species-level concordance. Raising identity thresholds to 98.7–99% removed roughly one third of NanoASV abundance and reduced species-level concordance. Hard identity filtering did not correct the residual gap or make full-length 16S quantitatively equivalent to WGS. These results do not validate every unrestricted best hit; ambiguity-aware multi-hit classification remains a separate question.

For both taxonomy and function, strong correlation of taxon or KO effect vectors does not imply equal effect magnitude, and significant multivariate age structure in both assays does not imply an identical developmental trajectory. The reproducibility of these broad signals supports their use in exploratory ecological screening, cohort stratification and hypothesis generation. For claims that depend on a specific species proportion, a specific KO abundance, accessory gene content or precise host-associated effect size, WGS remains the stronger measurement.

### WGS platform sensitivity

The use of two WGS platforms is a technical limitation. Sensitivity analyses indicate that it does not account for the main findings. Platform-associated differences were detectable for several RefSeq fine-rank correlation metrics, with MinION-associated WGS profiles showing lower 16S–WGS abundance correlation after age and kennel adjustment. Those effects were strongly attenuated under DogMAG, suggesting that the generic reference feature space is more sensitive to acquisition differences than the compact host-specific reference.

Platform adjustment had little effect on the rank ordering or magnitude of continuous age-signal transfer for either taxonomy or function. Paired assay-by-platform PERMANOVA interactions were small and non-significant, whereas age-by-assay interactions remained significant. Because PromethION and MinION were not randomized and the MinION branch contains proportionally more young animals, these estimates should not be interpreted as evidence for an intrinsic instrument effect. The reference, classifier and cross-assay developmental findings persisted after adjustment for WGS acquisition branch.

Several limitations remain. WGS is an operational comparator rather than biological ground truth, and the study lacks a mock community or orthogonal assay with known absolute composition. The RefSeq benchmark intentionally sacrifices reference breadth to enforce source-genome matching, whereas DogMAG is an in-cohort host-specific reference rather than an external validation set. PICRUSt2 predicts expected gene content from phylogeny and cannot recover sample-specific gene gain/loss or accessory elements; Kadath itself depends on assembly, gene prediction, annotation and read-back support. Repeated sampling is uneven, kennel and age are not experimentally balanced, and developmental analyses remain observational.

The matched design allowed these sources of variation to be examined separately. Both assays used the same DNA extracts from 97 specimens, with verified dog identity retained in repeated-measures analyses. We distinguished profile similarity from feature-level agreement and reproducibility of biological inference, held source genomes constant in the RefSeq benchmark, and compared classifiers within the same DogMAG reference. Analyses of multivariate age effects, feature detection and sequence-matching thresholds further identified where analytical harmonization reduced cross-assay differences and where discrepancies remained.

## Conclusions

Full-length ONT 16S and ONT WGS provide related but non-interchangeable descriptions of the canine gut microbiome. Across 97 matched specimens, 16S preserves broad functional profiles, much of the ecological structure among samples and a substantial fraction of the direction and ranking of age-associated biological change. Agreement is lower for individual-feature abundances and quantitative effect estimates.

Using the host-specific DogMAG reference improves genus/species agreement and age-signal transfer for both minitax and Kraken2, indicating that reference representation contributes substantially to discordance. Within each reference, minitax consistently yields stronger 16S–WGS concordance than Kraken2 across abundance, detection, ecological-distance and age-inference endpoints under both matched reference settings. Shared-feature restriction narrows the residual gap, whereas increasingly stringent NanoASV identity filtering does not.

Functional agreement is high for whole profiles and continuous age-associated KO effects, but lower for individual KO abundances; many KOs retain significant assay-by-age differences. Taxonomic PERMANOVA detects developmental structure in both assays, with a reproducible age-by-assay effect that is small relative to the overall assay difference.

Full-length ONT 16S is therefore suited to broad ecological screening, longitudinal community analysis and hypothesis generation. WGS remains preferable when conclusions require quantitative fine-rank composition, directly supported gene content, accessory-function detection or precise feature-level effect sizes.

## Declarations

### Ethics approval and consent to participate

In accordance with institutional and national regulations, formal animal ethics approval was not required for this study. The work involved only non-invasive collection of naturally voided feces during routine husbandry, without any change to housing, diet or veterinary care. Samples were provided voluntarily by the breeders, and all procedures complied with national animal welfare regulations.

### Availability of data and materials

The full-length ONT 16S source data used here are a subset of the longitudinal all-kennel dataset of Asaduzzaman et al. (Asaduzzaman et al. 2026) and are deposited in ENA under **PRJEB82125**. The matched WGS source reads span the relevant canine source projects **PRJEB82125** (DMD provenance), **PRJEB85420** (CaniMeta/Serteperti provenance) and **PRJEB115259** (new mixed-kennel WGS and DogMAG-associated records). DogMAG-derived reference resources and newly deposited genome/assembly records are associated with **PRJEB115259**. Analysis provenance and result locations are indexed in `documents/final_synthesis_manifest.tsv`. The project repository version-controls the canonical pairing, taxonomy-harmonization, NanoPredict integration and Kadath production scripts used for this comparison, including `scripts/run_kadath_single.sh` and `scripts/run_kadath_batch.sh`. A frozen archival release of the analysis code and final environment/version manifest should accompany submission. Minitax is available at https://github.com/Balays/minitax.

### Consent for publication

Not applicable.

## Authors’ contributions

N.J.Y. contributed to the investigation, DNA isolation, library preparation, and sequencing, and drafted the manuscript. B.K. analyzed the data, prepared visualizations, and wrote the manuscript. G.G. contributed to data handling and sequencing. M.A. contributed to DNA isolation and sequencing. Z.B. contributed to the scientific interpretation of the results and reviewed and edited the manuscript. D.T. conceived and supervised the study, led project administration, prepared visualizations, and contributed to drafting, reviewing, and editing the manuscript. All authors reviewed and approved the final manuscript.

## Acknowledgements

We would like to express our sincere gratitude to the following kennel owners for providing samples from their dogs: Ildikó Abonyi, owner of the Duna-menti Dumás Kennel (Baja, Hungary, https://www.pumikennel.eu/); Dr. Csaba Dobó-Nagy, owner of the Pattogó Parázs Kennel (Gödöllő, Hungary, https://pumi.pedigreedatabaseonline.com/hu/Pattog%C3%B3-Par%C3%A1zs-kennel-Csaba-Dobo-Nagy/breeder/638); Viktória Hordósné Vasas, owner of the Bükki Cserfes Pumi Kennel (Bükkzsérc, Hungary, https://www.xn--bkki-cserfes-pumi-kennel-vsc.hu); Gabriella Kassai, owner of the Serteperti Pumi Kennel (Kiskunmajsa [formerly Dány], Hungary, https://serteperti.hu/); Bianka Máthé-Szentes, owner of the Le Petit Lapin Pumi Kennel (Zomba, Hungary, https://pumi.mozellosite.com/) Melinda Takácsné Horváth, owner of the Rezerta-Réti Pumi Kennel (Ajka-Padragkút, Hungary, https://www.facebook.com/profile.php?id=100057382122448). Finally, we are especially grateful to Tünde Baloghné Jancsovics, a board member of the Hungarian Pumi Club (https://pumiklub.eu/), for establishing the crucial contact with breeders and facilitating the collaboration. We also thank Zsolt Csabai (DMB, USZ), Ákos Dörmő (DMB, USZ) and Tamás Járay (DMB USZ) for laboratory support and helpful discussions.

## Supplementary figures

**Supplementary Figure S1.**
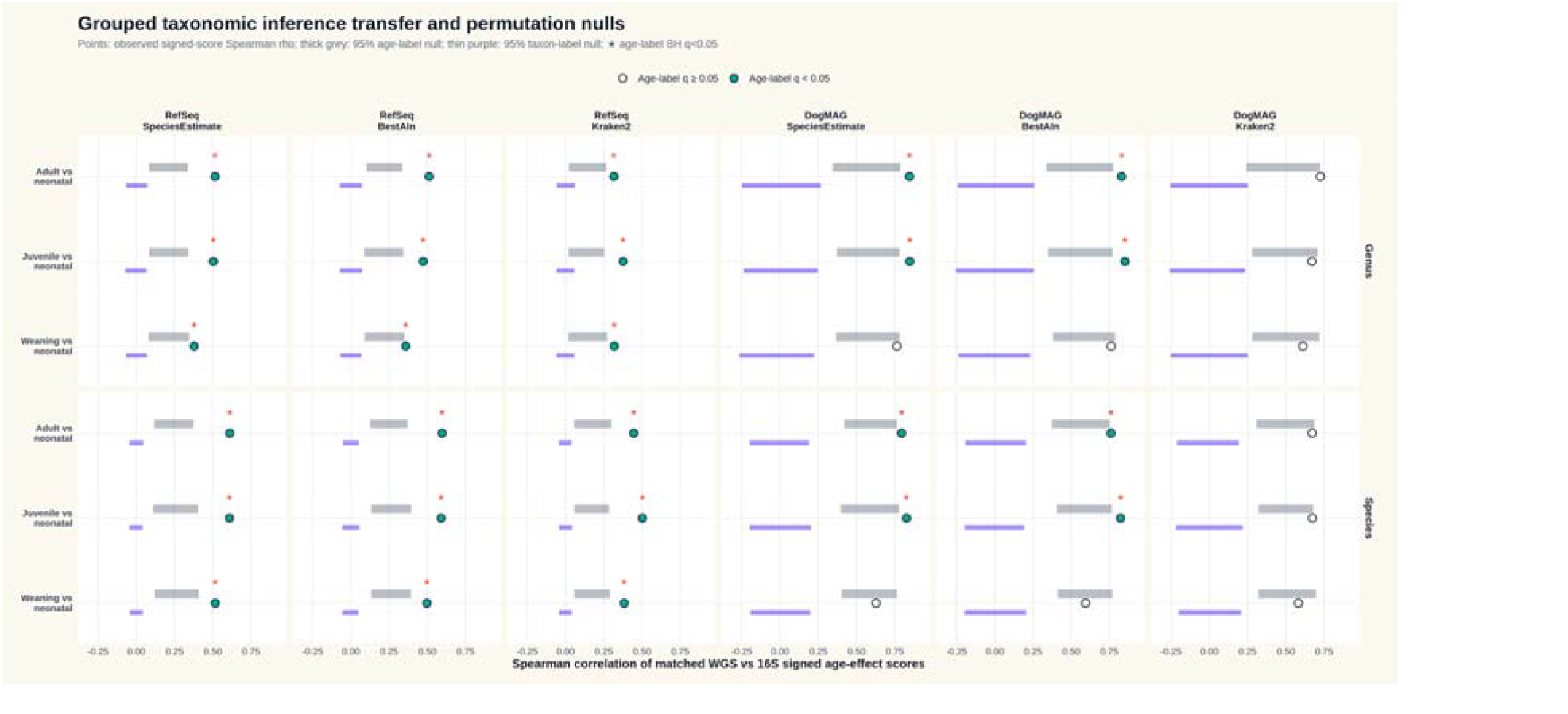
Grouped taxonomic inference transfer and permutation nulls. Observed Spearman correlations of matched WGS-versus-16S signed taxon age-effect scores are shown for each developmental contrast, taxonomic rank, reference context and classifier. Thick grey intervals show the 95% age-label permutation null; thin purple intervals show the 95% taxon-label permutation null. Filled points and stars indicate BH-adjusted age-label q<0.05. All displayed taxon-label permutation tests were significant at q=0.001. RefSeq shows significant age-label transfer across all grouped contrasts, DogMAG minitax shows robust juvenile/adult transfer but weaker earliest weaning transfer, and DogMAG Kraken2 retains high observed concordance without grouped age-label significance in the primary nonzero analysis.

## References

Aitchison, John. 1982. “The Statistical Analysis of Compositional Data.” Journal of the Royal Statistical Society: Series B 44: 139–77. 10.1111/j.2517-6161.1982.tb01195.x.

Albastaki, Abdulla, and Judith Smith. 2026. “Choosing Between Short-Read 16S, Full-Length ONT 16S, and Long-Read Shotgun Metagenomics for Soil Microbiome Studies: A Critical Review of the Benchmarking Evidence.” Microorganisms 14 (5): 1132. 10.3390/microorganisms14051132.

Asaduzzaman, Md, Péter Oláh, Natheer Jameel Yaseen, Ahmed Taifi, Tamás Járay, Gábor Gulyás, Zsolt Boldogkői, and Dóra Tombácz. 2026. “Longitudinal Long-Read Microbiome Profiling in a Canine Model Reveals How Age, Diet, and Birth Mode Shape Gut Community Dynamics.” mSystems 11 (2): e01279–25. 10.1128/msystems.01279-25.

Boldogkői, Zsolt, and Dóra Tombácz. 2026. “The Canine Gut Microbiome as a Translational Model for Human Health and Aging.” mBio, ahead of print. 10.1128/mbio.01951-26.

Cantalapiedra, Carlos P., Ana Hernández-Plaza, Ivica Letunic, Peer Bork, and Jaime Huerta-Cepas. 2021. “eggNOG-mapper v2: Functional Annotation, Orthology Assignments, and Domain Prediction at the Metagenomic Scale.” Molecular Biology and Evolution 38: 5825–29. 10.1093/molbev/msab293.

Castillo-Fernandez, Juan, Rachel Gilroy, Roshonda B. Jones, Ryan W. Honaker, Michaella J. Whittle, Phillip Watson, and Gregory C. A. Amos. 2026. “Waltham Catalogue for the Canine Gut Microbiome: A Complete Taxonomic and Functional Catalogue of the Canine Gut Microbiome Through Novel Metagenomic Based Genome Discovery.” Microbiome 14 (1): 25. 10.1186/s40168-025-02265-w.

Chitcharoen, Suwalak, Vorthon Sawaswong, Pavit Klomkliew, Prangwalai Chanchaem, and Sunchai Payungporn. 2025. “Comparative Analysis of Human Gut Bacterial Microbiota Between Shallow Shotgun Metagenomic Sequencing and Full-Length 16S rDNA Amplicon Sequencing.” BioScience Trends 19 (2): 232–42. 10.5582/bst.2024.01393.

Curry, Kevin D., Quang Wang, Michael G. Nute, Anna Tyshaieva, Elizabeth Reeves, Stephanie Soriano, Qinglong Wu, et al. 2022. “Emu: Species-Level Microbial Community Profiling of Full-Length 16S rRNA Oxford Nanopore Sequencing Data.” Nature Methods 19: 845–53. 10.1038/s41592-022-01520-4.

Cuscó, Anna, Carla Catozzi, Joana Viñes, Adrián Sanchez, and Olga Francino. 2018. “Microbiota Profiling with Long Amplicons Using Nanopore Sequencing: Full-Length 16S rRNA Gene and the 16S-ITS-23S of the Rrn Operon.” F1000Research 7: 1755. 10.12688/f1000research.16817.2.

Douglas, Gavin M., Vincent J. Maffei, Jesse R. Zaneveld, Svetlana N. Yurgel, James R. Brown, Christopher M. Taylor, Curtis Huttenhower, and Morgan G. I. Langille. 2020. “PICRUSt2 for Prediction of Metagenome Functions.” Nature Biotechnology 38: 685–88. 10.1038/s41587-020-0548-6.

Franzosa, Eric A., Lauren J. McIver, Gholamali Rahnavard, Levi R. Thompson, Melanie Schirmer, George Weingart, Karen S. Lipson, et al. 2018. “Species-Level Functional Profiling of Metagenomes and Metatranscriptomes.” Nature Methods 15: 962–68. 10.1038/s41592-018-0176-y.

Gulyás, Gábor, Balázs Kakuk, Ákos Dörmő, Tamás Járay, István Prazsák, Zsolt Csabai, Miksa Máté Henkrich et al. 2024. “Cross-Comparison of Gut Metagenomic Profiling Strategies.” Communications Biology 7: 1445. 10.1038/s42003-024-07158-6.

Hyatt, Doug, Gwo-Liang Chen, Philip F. LoCascio, Miriam L. Land, Frank W. Larimer, and Loren J. Hauser. 2010. “Prodigal: Prokaryotic Gene Recognition and Translation Initiation Site Identification.” BMC Bioinformatics 11: 119. 10.1186/1471-2105-11-119.

Járay, Tamás, Gábor Gulyás, Md Asaduzzaman, Ákos Dörmő, Zsolt Csabai, Balázs Kakuk, Zsolt Boldogkői, and Dóra Tombácz. 2026. “Early-Life Canine Gut Microbiome Maturation Follows a Shared Age–Diet Trajectory Within Persistent Host-Specific Structure.” bioRxiv. 10.64898/2026.05.25.727648.

Kakuk, Balázs, Ákos Dörmő, Ahmed Taifi, Tamás Járay, Gábor Kurucsai, Gábor Gulyás, István Prazsák, Zsolt Boldogkői, and Dóra Tombácz. 2026a. “Canine Fecal Microbiome Dataset: Ultra-Deep Multi-Platform Sequencing Across Extraction and Library Protocols.” Scientific Data 13: 1324. 10.1038/s41597-026-07594-5.

Kakuk, Balázs, Natheer Jameel Yaseen, Ákos Dörmő, Tamás Járay, Zsolt Boldogkői, and Dóra Tombácz. 2026b. “Dog-Wise Canine Gut Metagenome Assemblies with Reconstructed Bacterial Genomes and Viral Candidates.” bioRxiv. 10.64898/2026.09.01.747583.

Kanehisa, Minoru, Miho Furumichi, Yoko Sato, Mari Ishiguro-Watanabe, and Mao Tanabe. 2021. “KEGG: Integrating Viruses and Cellular Organisms.” Nucleic Acids Research 49 (D1): D545–51. 10.1093/nar/gkaa970.

Kolmogorov, Mikhail, Jeffrey Yuan, Yu Lin, and Pavel A. Pevzner. 2019. “Assembly of Long, Error-Prone Reads Using Repeat Graphs.” Nature Biotechnology 37: 540–46. 10.1038/s41587-019-0072-8.

Li, Heng. 2018. “Minimap2: Pairwise Alignment for Nucleotide Sequences.” Bioinformatics 34: 3094–3100. 10.1093/bioinformatics/bty191.

Mantel, Nathan. 1967. “The Detection of Disease Clustering and a Generalized Regression Approach.” Cancer Research 27: 209–20.

Matchado, Monica Steffi, Malte Rühlemann, Sandra Reitmeier, Tim Kacprowski, Fabian Frost, Dirk Haller, Jan Baumbach, and Markus List. 2024. “On the Limits of 16S rRNA Gene-Based Metagenome Prediction and Functional Profiling.” Microbial Genomics 10 (2): 001203. 10.1099/mgen.0.001203.

Ni, Yuxin et al. 2023. “The Newest Oxford Nanopore R10.4.1 Full-Length 16S rRNA Sequencing Enables the Accurate Resolution of Species-Level Microbial Community Profiling.” Applied and Environmental Microbiology 89: e00605–23. 10.1128/aem.00605-23.

Polacchini, Giulia, Bruno Stefanon, Paolo Mongillo, and Danilo Licastro. 2026. “A Comparative Analysis of Gut Microbiome in Dogs Using Short- and Long-Reads of 16S rRNA Sequences Reveals Workflow-Dependent Biases.” Veterinary Research Communications 50 (5): 463. 10.1007/s11259-026-11407-w.

Sun, Shan, Roshonda B. Jones, and Anthony A. Fodor. 2020. “Inference-Based Accuracy of Metagenome Prediction Tools Varies Across Sample Types and Functional Categories.” Microbiome 8: 46. 10.1186/s40168-020-00815-y.

Tamames, Javier, and Fernando Puente-Sánchez. 2019. “SqueezeMeta, a Highly Portable, Fully Automatic Metagenomic Analysis Pipeline.” Frontiers in Microbiology 9: 3349. 10.3389/fmicb.2018.03349.

Wood, Derrick E., Jennifer Lu, and Ben Langmead. 2019. “Improved Metagenomic Analysis with Kraken 2.” Genome Biology 20: 257. 10.1186/s13059-019-1891-0.

